# PHAIR – A biosensor for pH measurement in air-liquid interface

**DOI:** 10.1101/2020.11.09.375683

**Authors:** Mohammadhossein Dabaghi, Neda Saraei, Gang Xu, Abiram Chandiramohan, Jonas Yeung, Jenny P. Nguyen, Milica Vukmirovic, Ponnambalam Ravi Selvaganapathy, Jeremy A. Hirota

**Affiliations:** Firestone Institute for Respiratory Health–Division of Respirology, Department of Medicine, McMaster University, Hamilton, ON, L8N 4A6, Canada; Department of Mechanical Engineering, McMaster University, Hamilton, ON, L8S 4L7, Canada; School of Biomedical Engineering, McMaster University, Hamilton, ON, L8S 4K1, Canada; McMaster Immunology Research Centre, Department of Pathology and Molecular Medicine, McMaster University, Hamilton, ON, L8S 4K1, Canada; Division of Respiratory Medicine, Department of Medicine, University of British Columbia, Vancouver, BC, V6H 3Z6, Canada; Department of Biology, University of Waterloo, Waterloo, ON, N2L 3G1, Canada

**Keywords:** pH, Wire-format, air-liquid interface, cell culture, airway surface lining, epithelial

## Abstract

In many biological systems, pH can be used as a parameter to understand and study cell dynamics. However, measuring pH in live cell culture is limited by the sensor ion specificity, proximity to the cell surface, and scalability. Commercially available pH sensors are difficult to integrate into a small-scale cell culture system due to their size and are not cost-effective for disposable use. We made PHAIR - a new pH sensor that uses a micro-wire format to measure pH in vitro human airway cell culture. Tungsten micro-wires were used as the working electrodes, and silver micro-wires with a silver/silver chloride coating were used as a pseudo reference electrode. pH sensitivity, in a wide and narrow range, and stability of these sensors were tested in common standard buffer solutions as well as in culture media of human airway epithelial cells grown at the air-liquid interface in a 24 well cell culture plate. When measuring the pH of cells grown under basal and challenging conditions using PHAIR, cell viability and cytokine responses were not affected. Our results confirm that micro-wires-based sensors have the capacity for miniaturization, and detection of diverse ions while maintaining sensitivity. This suggests the broad application of PHAIR in various biological experimental settings.

## 2 Introduction

Changes in extracellular or intracellular pH may accompany cellular growth, metabolism, signaling, or ion transport ^1,2^. For instance, as cancer cells proliferate, they produce lactic acid that reduces extracellular pH in the local microenvironment ^2,3^. The central role that pH plays in cell biology highlights the importance of dynamic measurement of extracellular pH in cell culture experiments in both discovery and applied biomedical research, under basal and challenging conditions. ^4–6^. Human airway epithelial cells (HAECs) are an indispensable component of the innate immune system functioning as the first line of defense in the lungs, protecting against inhaled pathogens and environmental toxins, which may imbalance the extracellular pH of cells ^7^. In addition to serving as physical barriers, HAECs secrete a thin layer of airway surface lining (ASL) fluid that functions to hydrate the airways, facilitate mucociliary clearance, and trap inhaled substances. ASL, while mostly composed of water, contains salts, lipids, mucus, and protective proteins, including antimicrobial factors ^8,9^. Recent evidence suggests that the antimicrobial activities exhibited by ASL and some of its constituents, such as lactoperoxidase, LL-37, beta-defensin 1 (hBD-1), and beta-defensin 3 (hBD-3) are pH-dependent ^8,9^. In addition, ASL pH has been shown to modulate other aspects of airway homeostasis, ranging from altering mucus properties to regulating ionic movements ^9,10^.

ASL pH is regulated by a variety of mechanisms involving paracellular pathways and membrane transport proteins, among which, Cystic Fibrosis Transmembrane Conductance Regulator (CFTR – also known as ABCC7) has received emphasis as the genetic defect behind Cystic Fibrosis (CF). CFTR activity is potentiated by cyclic-AMP (cAMP)-dependent protein kinase A phosphorylation of the intracellular regulatory region ^11^. Classically thought to be a Cl^-^ channel, CFTR also conducts HCO_3_^-^ and has been identified as a crucial mechanism of base secretion in the airways ^9,12,13^. In cellular and animal models of CF, ASL acidification resulting from defective CFTR function has been demonstrated, leading to the hypothesis that abnormal ASL pH may reduce antimicrobial activity and contribute to chronic bacterial infections, a characteristic feature of CF airway disease ^14–20^. Accordingly, pharmacological interventions that aim to normalize ASL pH by increasing CFTR activity or targeting other acid/base transport proteins have been investigated ^20,21^. Although the lack of specificity of forskolin, a cAMP-elevating agent, renders it unsuited for clinical use, it serves as a robust tool for inducing CFTR *in vitro* research and drug response assays ^22^. Forskolin directly activates adenylate cyclase, resulting in an increase in intracellular cAMP concentration. Through a signaling cascade, phosphorylation-dependent CFTR modulation occurs, leading to potentiation of activity and ion flux (Li et al., 2005; Schnur et al., 2019).

Performing simple yet accurate and reliable measurements of ASL pH has posed a significant challenge, with a variety of methods deployed to date ^14,15,17,19–21,25,26^. A particular challenge exists for pH measurements of airway epithelial cells grown under air-liquid interface culture conditions on Transwell inserts as there is a small growth surface area contributing to a low volume and depth of ASL. Since the measurement of pH in a cell culture system has been recognized as a useful tool to study different aspects of the cells’ biology, various pH sensors have been developed ^27–32^. Several techniques, such as optical sensing methods (Magnusson et al., 2013; Munoz-Berbel et al., 2013; Shaegh et al., 2016), ion-sensitive field-effect transistor-based sensors, or electrochemical pH sensing materials ^36^ have been investigated to monitor pH in a biological application. Among these methods, electrochemical-based pH sensors have great potential to be miniaturized and integrated into a cell culture system, especially potentiometric sensors. It is well-known that various metal oxides such as copper oxide ^36^, iridium oxide ^5,37–40^, tungsten oxide ^41–44^, cobalt oxide ^45^, titanium oxide ^45^, ruthenium oxide ^39^, zinc oxide ^46^, indium tin oxide ^47–49^, and palladium oxide ^50–53^ can be used as a pH sensing material in a potentiometric sensor. The fabrication of this type of pH sensor can be simplified to develop a straightforward compact pH sensor that can be easily integrated into different cell culture systems. Of the candidate materials, tungsten is always oxidized at room temperature resulting in a surface oxide layer with sensitivity to changes in pH ^41,42^. Based on the above, we hypothesized that a tungsten wire-based pH sensor system satisfies design constraints for pH measurements in airway surface lining fluid from human airway epithelial cells cultured under air-liquid interface conditions.

Unlike other metallic oxides, tungsten is a metal that is oxidized at room temperature under a dry atmosphere resulting in the formation of a thin layer of WO_3_ on the tungsten surface ^41^. Exposing tungsten to an aqueous environment also results in the formation of a thicker layer of WO_2_, WO_3_, and hydroxide ^41^. When tungsten is placed in a buffer with a pH around 7, a slightly different form of tungsten oxides (WO_2_ or W_2_O_5_) can be formed ^42^. The formed metal oxide layer is responsible for the pH response of tungsten. Measuring the open circuit potentials (E) between the working electrode (metal/metal oxide) and a reference electrode, which are placed in an aqueous solution, can reveal information about the pH of the solution of interest. The metal/metal oxide electrode and its aqueous surrounding exchange proton (H^+^) and this redox reaction will finally reach an equilibrium at a constant pH, as shown below ^44^:

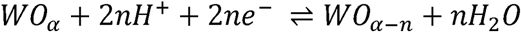

In this work, we developed pH sensors using tungsten wire and demonstrated the capacity to measure pH in vitro cell culture. In order to miniaturize the sensor, we first developed a custom-made silver/silver chloride (Ag/AgCl) reference electrode in a micro-wire format to be able to deploy in HAEC ASL fluid for *in situ* measurement of pH. Secondly, the pH sensitivity of tungsten wires was studied relative to a commercial reference electrode or the optimized custom-made silver/silver chloride reference electrodes. Thirdly, we evaluated our pH sensor in different buffer standard solutions, non-buffer solutions, and cell culture media to assure that the pH sensor was robust and maintained sensitivity.

## 3 Experimental

### 3.1 Electrode preparation and fabrication of reference electrodes

Tungsten (W) wires (99.95% metals basis) and silver (Ag) wires (99.9% metals basis) were purchased from Alfa Aesar with a diameter of 250 µm, cut into 10-cm long pieces. The wires were stored at room temperature (RT) prior to preparation for each experiment. Tungsten wires acted as a working electrode (WE) to measure pH. Three fabrication processes were used to make four custom-made silver/silver chloride (Ag/AgCl) reference electrodes (REs) as shown in Scheme 1) silver wires were immersed in silver/silver chloride ink or paste (ratio of 65:35, product # 125-21, Creative Materials, Ayer, MA, USA) for 5 minutes (Scheme 1a1), wiped by a disposable swab to remove extra ink and achieve a thin layer of ink on wires (Scheme 1a2), dried in an oven at 65 °C overnight, named as custom-made RE1 (Scheme 1a3) Silver wires were immersed in a bleach solution (Javex, Fisher Scientific, Canada) containing 10.8 % sodium hypochlorite (NaOCl) or NaOCl solution (product # 425044, Sigma Aldrich) containing 10 – 15 % chlorine for 5 minutes (custom-made RE2) or 10 minutes (custom-made RE3) as seen in Scheme 1b1, rinsed rigorously with deionized water (DI), and dried at RT (Scheme 1b2). 3) Silver wires were immersed in bleach or NaOCl solution for 5 minutes (Scheme 1c1), rinsed with DI water, dried at RT overnight (Scheme 1c2), immersed in silver/silver chloride ink for 5 minutes (Scheme 1c3), wiped by a disposable swab (Scheme 1c4), and dried in an oven at 65 °C overnight, named as custom-made RE4 (Scheme 1c5).

**Scheme 1:**
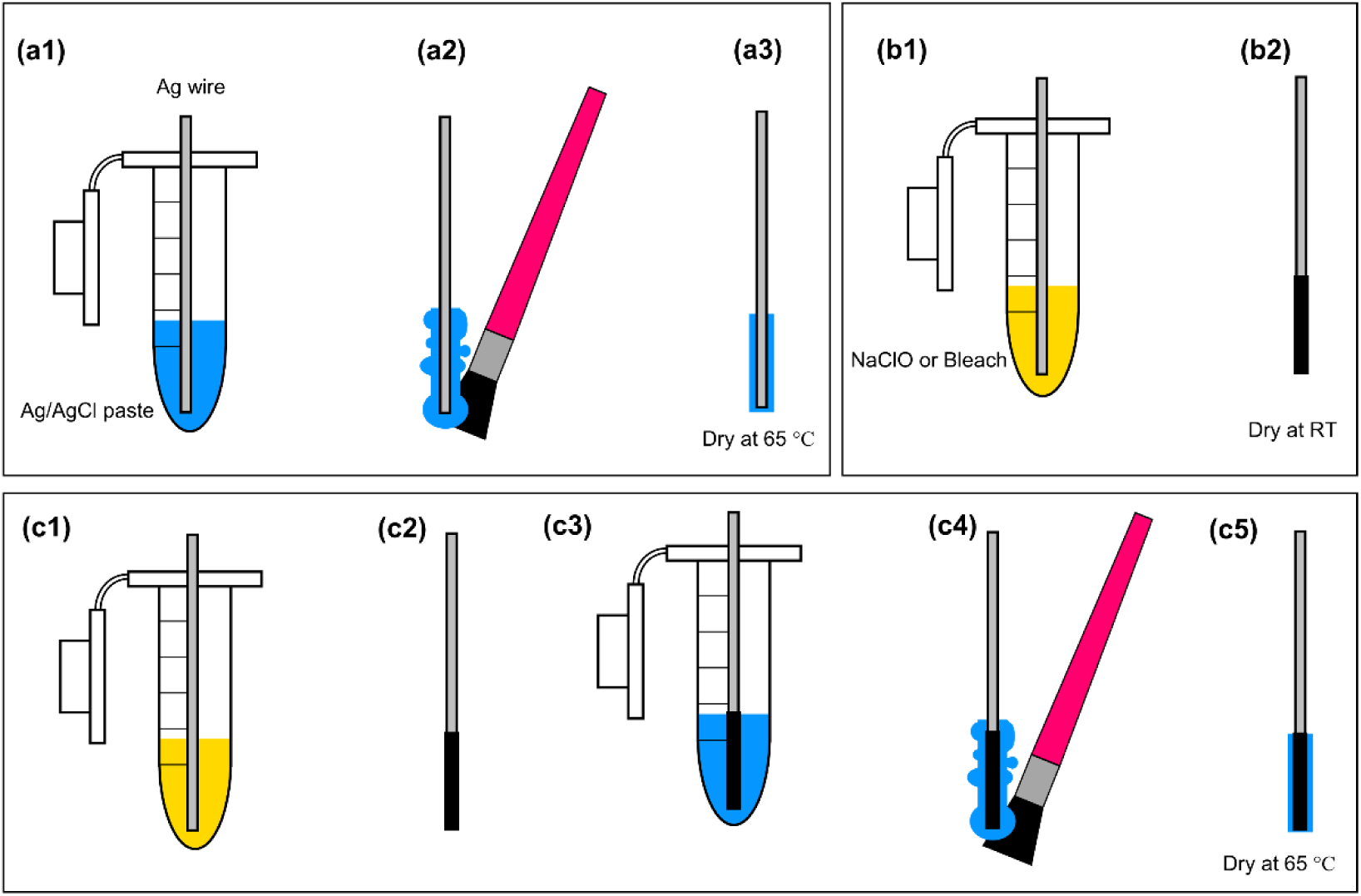
The fabrication process of miniaturized reference electrodes in a wire format: (a1 – a3) a 10-cm long silver wire (diameter = 250 µm) was immersed in silver/silver chloride paste for 5 minutes, extra paste on the wire was removed by a disposable brush, and the coated silver wire was cured in an oven at 65 °C overnight, (b1 – b2) a 10-cm long silver wire was immersed in a commercial bleach or sodium hypochlorite (NaClO) solution for 5 or 10 minutes, rinsed rigorously with DI water, and dried at RT, and (c1 – C5) a 10-cm long silver wire was first chlorinated in a commercial bleach solution of NaClO for 5 minutes, rinsed with DI water, dried in an oven at 65 °C for 2 – 3 hours, immersed in silver/silver chloride paste for 5 minutes, extra material was removed and dried in in an oven at 65 °C.

### 3.2 Electrochemical sensing tests

For all electrochemical experiments, a QUAD ISOPOD e-DAQ system was used to measure open circuit potentials (E) of tungsten wires with respect to a commercially-available reference electrode (CRE) or one of the custom-made reference electrodes. The commercial reference electrodes were a miniature silver/silver chloride reference electrode that had a porous Teflon tip, and they were purchased from CH Instruments, Inc. (CHI111, Austin, USA). The commercial reference electrodes were stored in a 3M potassium chloride (KCl) solution at RT. To analyze the pH response of tungsten wires against a commercial reference electrode or one of the custom-made reference electrodes, a set of wires consisted of a tungsten wire, and one of the reference electrodes was immersed in a buffer solution with known pH. Immediately after immersion, E was recorded for 5 minutes, the recording was paused, the wire setup was removed from the buffer solution, rinsed with DI water, transferred to the next buffer solution, followed by the continuation of recording with 10-second delay to let wires wet properly in the new buffer. Three pH buffer standard solutions with pH of 4, 7, and 10 were used to test the pH sensitivity and stability of tungsten wires against a commercial reference electrode after one day and four days storage in phosphate-buffered saline (PBS) with a pH of ∼ 7.4. The standard buffer solution with a pH of 10 was used for measuring the response time of each sensor setup. Hydrion buffers from Micro Essential Laboratory, Inc. with various pH (pH = 7, 8, and 9) were also used for testing the sensors for smaller pH intervals. The pH of these buffers was adjusted to the desired value if needed by adding 1 M sodium hydroxide (NaOH). To test and investigate the pH sensitivity of the sensors in a complex environment, Medium 199 (M0393, Sigma Aldrich) containing Hanks′ salts and L-glutamine without sodium bicarbonate was used. After dissolving the medium powder in DI water, the pH of the medium was adjusted to ∼ 6.75, 7.50, 8.25, and 9.00 by adding 1 M NaOH (the medium was acidic with a pH of ∼ 6.00). All experiments were performed at room temperature. The pH of each media was measured before and after each step to detect any pH drift, since these media did not contain a buffering system, and no significant change in pH was observed.

### 3.3 Fabrication and assembly of macro-wells with wire sensors

Polydimethylsiloxane (PDMS) monomer base and its curing agent were mixed at a ratio of 10:1 and degassed in a desiccator for 15 minutes. PDMS was poured to a petri dish (thickness 8 mm) and cured in an oven at 65 °C for 2 hours. A disposable punch (Accu-Punch 12.0 mm, Electron Microscopy Sciences, PA, USA) with a diameter of 12 mm was used to punch PDMS slabs and create macro-wells (Scheme 2a). Tungsten wires were pushed through from the sides and inserted in the wells (Scheme 2b). To insert the custom-made reference electrode into each well, a six mm-deep cut was made from the top of each well by a scalpel (Scheme 2c). To seal the bottom of each well, the bottom side of wells and another cured PDMS piece were treated by a butane torch (Scheme 2c), and a custom-made reference electrode was carefully placed inside the bottom of the created slot of each well (Scheme 2d). Then, wells with inserted wires and another PDMS piece were brought into contact, followed by a gentle press to initiate bonding. Uncured PDMS was used to seal the cuts, and the product was placed in an oven at 65 °C overnight to strengthen the bonding (Scheme 2d).

**Scheme 2:**
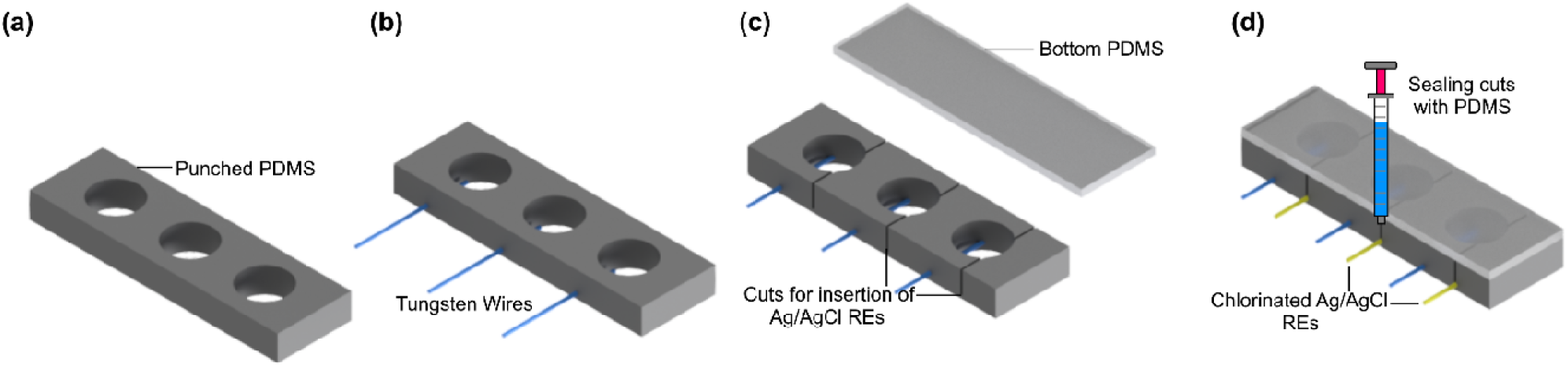
The fabrication process of the macro-well array: (a) the PDMS was cut and punched to create wells, (b) tungsten wires were pushed through PDMS to be placed into wells, (c) a cut (1 mm from the bottom) was made followed by treating flame activation for bonding, and (d) silver/silver chloride reference electrodes were placed in their locations and two pieces were brought in contact to initiate the bonding followed by sealing cuts using PDMS.

### 3.4 Cell culture

The human airway epithelial cell line, Calu-3 (HTB-55™, ATCC®, USA), was used for testing the effect of wires on cell viability and cytokine release. The cells were cultured in alpha minimum essential medium (αMEM, Corning®, USA) which was supplemented with fetal bovine serum (10%, FBS, WISENT Inc., Canada), and antibiotic-antimycotic (100 U/mL penicillin, 100 µg/mL streptomycin, 0.25 µg/mL amphotericin B, Gibco®, USA) and HEPES buffer (10 µM, Gibco®, USA). The cells were cultured and maintained in an incubator at 37 °C with 5% carbon dioxide, and they were fed on a two-day feeding cycle.

Prior to seeding cells in PDMS wells, the bottom of wells was coated with type I collagen solution (PureCol®, Bovine Collagen, 3 mg/mL, Advanced BioMatrix, CA, USA). The collagen solution had a concentration of 3 mg/mL and was diluted to 1 mg/mL with autoclaved DI water. To coat the wells, 200 µL of 1 mg/mL collagen solution was added to each well, and all wells were stored in the fridge for 24 hours. Then, the collagen solution was removed from wells, wells were rinsed with PBS (pH = 7.4) for at least three times, the wells were dried in a biosafety cabinet, and they were sterilized by UV light for 30 minutes. Calu-3 cells with a density of 10 × 10^5^ cells/well were cultured in the wells and were grown to confluency for viability assay. Calcein AM (5 µM, Life Technologies®), and Hoechst (20 µg/mL, Thermo Fisher Scientific, USA) dyes were used to perform quantitative viability assay. First, the medium was removed from the wells, at the cells were gently washed with warmed PBS two times, 400 µL of Calcein AM and Hoechst solution was added to each well, they were incubated with dyes at 37 °C for 20 minutes, the dyes were removed, they were rinsed with warmed PBS for at least two times, and they were imaged by an EVOS M7000 microscope (Thermo Fisher, Canada). Wells without wires were also fabricated, cultured with Calu-3 cells, and imaged for controls.

### 3.5 Enzyme-Linked Immunosorbent Assay (ELISA) and Cell viability assessment

Cell culture media was collected and spun down at 7500 xg for 15 minutes at 4 °C. Supernatants were subsequently analyzed for IL-6 and IL-8 cytokines via ELISA. Human IL-6 Duoset (R&D Systems; Catalog Number DY206) and Human IL-8/CXCL8 Duoset (R&D Systems; Catalog Number DY208) were used in accordance with the manufacturer’s protocols. The following maximum and minimum limits of standards were used, respectively: IL-6 (1200 pg/ml, 0 pg/ml) and IL-8 (4000 pg/ml, 0 pg/ml) with the assay plate analyzed as per manufacturer’s protocols using a SpectraMax i3x microplate reader.

Cell viability was determined using secreted lactate dehydrogenase (LDH) using a commercially available kit (Thermofisher; Catalog Number C20301) and following manufacturer’s directions.

Calu-3 cells were cultured in a 24-well plate with a density of 10 × 10^5^ cells/well and grown to confluency. Then, a set of two-wire sensing system including a 10 mm-long tungsten wire and a 10 mm-long custom-made reference electrode (5 minutes bleached silver wires) was transferred to four wells, and the cells were maintained for two days. Supernatants were collected for ELISA and LDH assays. Positive controls indicating maximal LDH release were generated by seeding a 24-well cell culture plate with the same cell density and incubating in 1 mL of cell culture media until confluency followed by the addition of 100 µL of lysis buffer provided in the assay kit as per manufacturer’s directions. All supernatants were then harvested and spun down at 7500 xg for 15 minutes before use in LDH and ELISA assays.

### 3.6 Calu-3 Cell Culture at Air-Liquid Interface

Calu-3 cells were seeded onto semi-permeable Transwell insert (Corning) at a density of 100,000 cells/well, with 200 µL and 750 µL of media in the apical and basal compartments, respectively. Media were replaced every two days. When confluency was reached after five days, the apical medium was removed, and cells were fed basal-only every two days to allow for differentiation at the air-liquid interface (ALI day 0). In addition, transepithelial electrical resistance (TEER) measurements were performed as quality control after feeding by adding 200 µL of Hanks’ Balanced Salt Solution (HBSS) to the apical compartment and using the TEER machine (Millicell® ERS-2 Voltohmeter) according to the manufacturer’s protocol. After TEER measurements, the apical surface of cells was gently washed with 200 µL of HBSS for 10 min to remove debris and mucus. On ALI day 16, one hour prior to stimulation, the basal medium was replaced, three apical washes with HBSS were performed, and TEER values were >700 Ω·cm^2^ for all wells.

### 3.7 Forskolin stimulation and airway surface liquid pH Measurements

To allow for measurement of ASL pH changes, 100 µL of HCO_3_^-^ and K^+^-free saline Ringer’s solution (140 mM Na^+^/140 mM Cl^-^/1.2 mM Ca^2+^/1.2 mM Mg^2+^/2.5 mM PO_4_ ^3-^, pH 6) was added to the apical compartment of Transwell inserts. After the addition of 10 µM of forskolin or dimethyl sulfoxide (DMSO) vehicle control to the basal medium, cells were incubated for 3 hours in an incubator and subsequently left at room temperature for 30 minutes. ASL pH for each well was then measured using our pH sensor, followed immediately by a commercial pH microelectrode (Orion™ PerpHecT™ ROSS™). Sensor fabrication and setup were carried out in accordance with the previously described protocol. Calibration was performed in HCO_3_^-^ and K^+^-free saline Ringer’s solutions (pH 6, 6.75, 7.5, 8.25, and 9) at room temperature to obtain ascending and descending calibration curves. The sensor wires were conditioned in the pH 6 solution for at least 2 hours prior to use. To simplify the measurement process and ensure wire stability in solution, a custom holder was designed to fit onto Transwell inserts and was 3D-printed using A FormLab Form 2 printer (MA, USA); the working and reference electrodes were firmly attached to the 3D-printed holder with instant glue (Loctite 4013). For each well, open circuit potential (mV) was recorded every second for a total duration of 150 seconds. The last 60 values were averaged, and corresponding pH values were obtained from the ascending and descending calibration curves. The apical solution was subsequently transferred from the Transwell insert into a micro Eppendorf tube, allowing for pH measurement with a commercial pH meter. At the end of the experiment, live and dead cell viability assay from ThermoFisher was used to assess the viability of cells.

## 4 Results and discussion

### 4.1 Tungsten wire pH response and stability against a commercial reference electrode

In this work, the potentiometric measurements of the metallic tungsten wires were conducted in a series of standard buffer solutions with a pH of 10, 7, and 4. Initially, tungsten wires were placed in the pH 10 standard buffer solution, and E was continuously recorded for 20 minutes. The aim of the experiment was to evaluate the pH response time of tungsten wires when they were transferred from a dry condition (room air) to an aqueous environment. In this study, the response time of tungsten wires is defined as the time that the sensors need to reach 90 % or 95 % of their equilibrium potential ^40^. Using this definition, tungsten wires reached 90 % and 95 % of their equilibrium potential in 12 seconds and 58 seconds, respectively, as presented in Figure 1a. A change in potential from ∼ -500 mV to ∼ - 432 mV was observed, attributable to the transformation of the oxide layer from WO_2_ to WO_2_/WO_3_/W_2_O_5_ ^41,42^. Then, tungsten wires were stored in PBS with a neutral pH for one day before performing pH sensitivity testing. One-day storage in a buffer solution would ensure that all tungsten wires were uniformly oxidized. Figure 1b shows the real-time pH response of the metallic tungsten wires against a commercial reference electrode. A smooth transition of the potentials was observed while the tungsten wires were transferred from one buffer to another one. Figure 1c to Figure 1f show measured E of the tungsten wires versus pH (the calibration curve for each sensor), where all of them demonstrated a good linear property (R^2^ > 0.999). Moreover, the hysteresis between the measured open circuit potential for descending and ascending pH was small and negligible (< 1%; red lines and black lines had covered each other).

**Figure 1:**
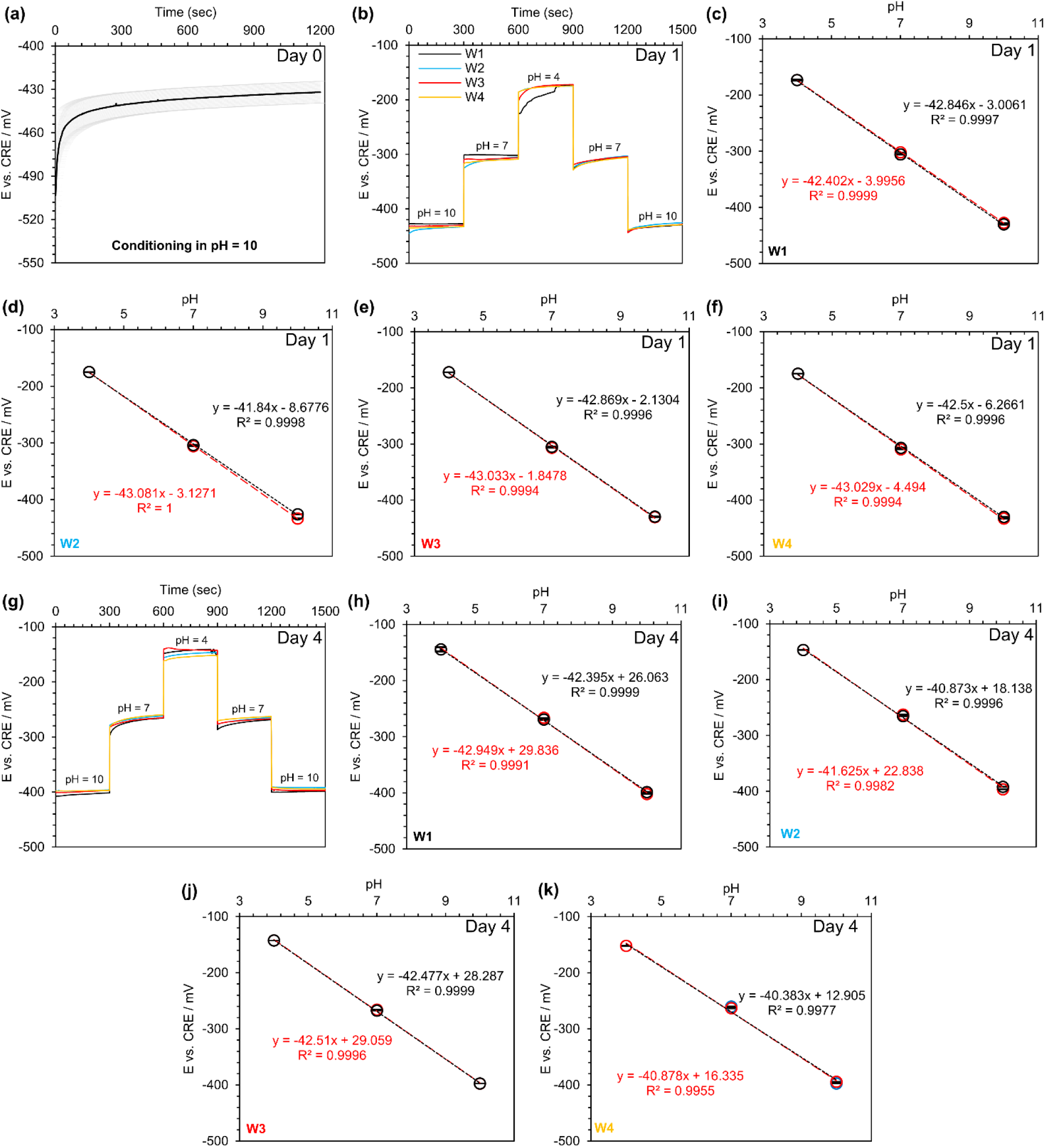
Open circuit potential (E) versus a commercial reference electrode (CRE) for tungsten (W) wires: (a) the stability testing of tungsten wires (n=7) over 20 minutes in a buffer with the pH of 10, (b) real-time response of tungsten wires in 3 standard pH buffers (4, 7, and 10) after being conditioned in PBS (pH = 7.4) for one day, (c – f) the calibration curve for each individual wire at day one, (g) real-time response of tungsten wires in 3 standard pH buffers (4, 7, and 10) after being stored in PBS (pH = 7.4) for four days, and (h – k) the calibration curve for each individual wire at day four. N = 4. The decrease in pH is shown by red colors, and the increase in pH is shown in black color.

One of the main challenges for any type of pH sensor is its stability over time. Here, the tungsten wires were tested on day 4 again under identical conditions, and their pH behavior was studied at day four (Figure 1g). Compared to day 1, measured E was changed in a range of 7 % to 15 % (∼ 7 % for the basic buffer, ∼ 13 % for the neutral buffer, and ∼ 15 % for the acidic buffer). This behavior suggested that longer storage in an aqueous solution would change the oxidized layer properties (the combination of metal/metal oxide or the thickness of metal oxide layer) as observed in other studies as well ^41,42^. However, the tungsten wires still exhibited a linear response (R^2^ > 0.995), and the slope did not change significantly compared to day 1 (the average slopes at day 1 and day 4 were – 42.7 ± 0.5 mV/pH and – 41.8 ± 1.0 mV/pH, respectively) as depicted in Figure 1g-k.

### 4.2 Characterization of custom-made reference electrodes

The pH sensitivity of metallic tungsten facilitated the fabrication of a miniature pH sensor in a micro-wire format with tungsten metal forming the basis of the working electrode. Nonetheless, a potentiometric pH sensor consisted of a working electrode and a reference electrode. The majority of commercially-available reference electrodes have a macro size (typically they have a diameter greater than 1 - 5 mm) and cannot be integrated into a micro-scale pH sensor device. As a result, a micro-scale reference electrode was required to realize a miniaturized pH sensor. Silver wires with a diameter of 250 µm were used as a substrate to fabricate four types of silver/silver chloride reference electrodes. For each electrode type, six reference electrodes were prepared, immersed in a standard buffer with a pH of 10, and their real-time response against a tungsten wire was monitored over 30 minutes, as seen in Figure 2. All custom-made reference electrodes were air-dried and used fabrication immediately. Custom-made RE1 had the largest variation, and the equilibrium potential varied from – 625 mV to – 586 mV (Figure 2a). Chlorinated REs (RE2 and RE3) had the smallest variation for the equilibrium potential ranging from – 661 mV to – 641 mV (Figure 2b and Figure 2c). The equilibrium potential of custom-made RE4 was also as small as chlorinated reference electrodes. However, custom-made RE4 had a noisier response compared to chlorinated reference electrodes (Figure 2d). Therefore, the chlorination of silver wires for 5 minutes was chosen as the standard process for the fabrication of all reference electrodes in this study.

**Figure 2:**
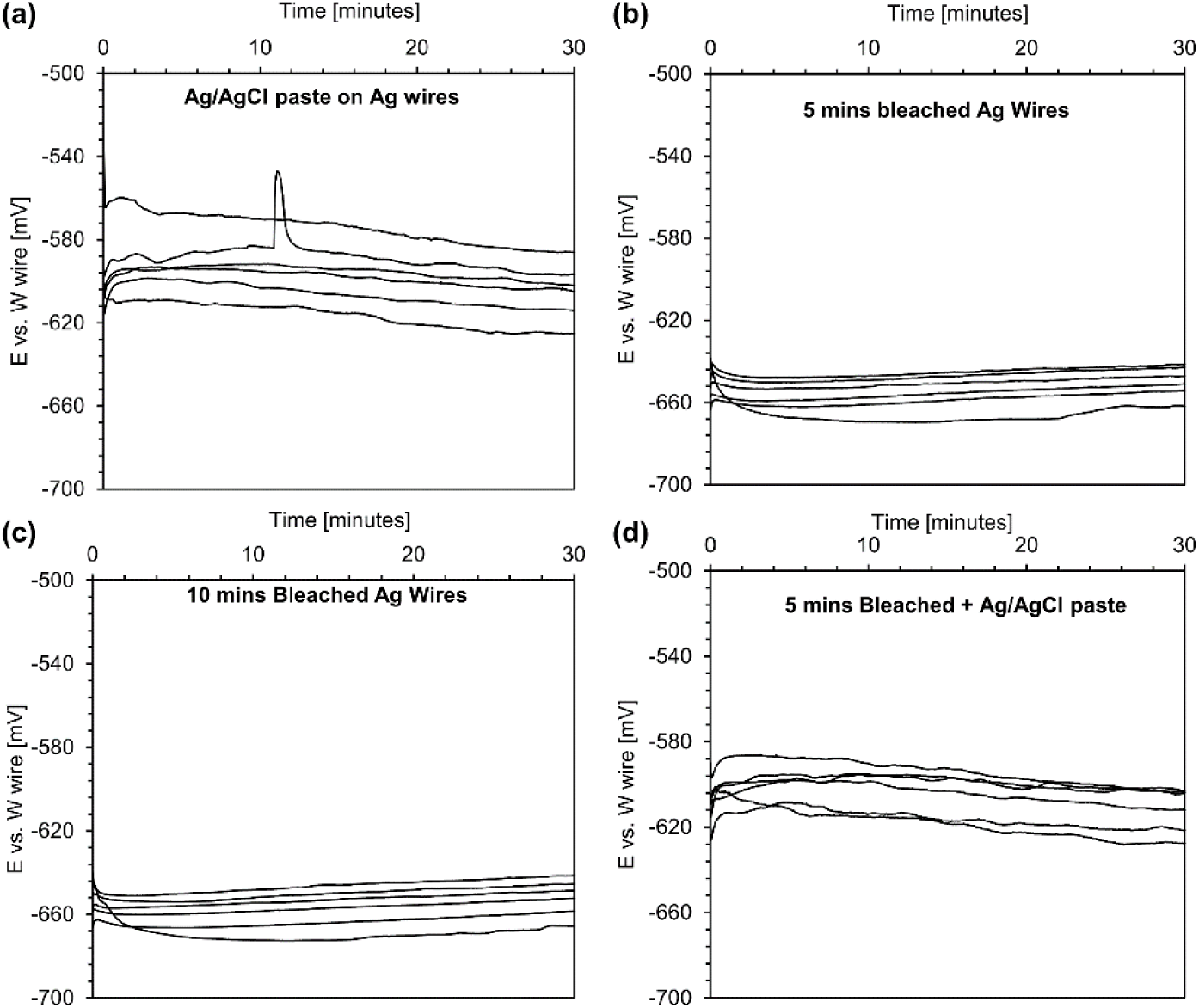
The stability evolution of custom-made reference electrodes: (a) real-time response of silver wires coated with silver/silver chloride paste versus a tungsten wire in a buffer with the pH of 10 for 30 minutes-RE1, (b) real-time response of 5-minutes bleached silver wires versus a tungsten wire in a buffer with the pH of 10 for 30 minutes-RE2, (c) real-time response of 10-minutes bleached silver wires versus a tungsten wire in a buffer with the pH of 10 for 30 minutes-RE3, and (d) real-time response of 5-minutes bleached silver wires + silver/silver chloride paste versus a tungsten wire in a buffer with the pH of 10 for 30 minutes-RE4. N = 6.

The main drawback of applying silver/silver chloride paste was that commercially available inks had supplier dependent properties and batch to batch variations, which made the replication of the results difficult. The chlorination of silver wires was recognized as a versatile method for producing silver/silver chloride reference electrodes and was less dependent on a given chlorine solution. Therefore, it was realized that reference electrodes chlorinated by NaClO solution were more stable and showed less variation in their potentials. Immersion of silver wires in a bleach solution or NaClO solution (10 – 15 %) led to a chemical reaction at the surface of silver wires as follows ^54^:

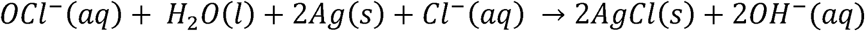

### 4.3 pH response and stability: Combination of tungsten wire and chlorinated silver wire reference electrodes

The silver wires chlorinated for 5 minutes were conditioned in 3M KCl, PBS, and room air, and their real-time response was studied against a tungsten wire in different standard pH buffers (Figure 3a). Chlorinated silver wires that were conditioned in PBS showed a more stable response and less drifting compared to other conditions. The pH sensitivity and real-time response of custom-made reference electrodes that were conditioned in 3M KCl continuously drifted relative to dry, and PBS conditioned electrodes. Consequently, for the rest of the experiments, custom-made reference electrodes were conditioned and stored in neutral PBS unless otherwise mentioned.

**Figure 3:**
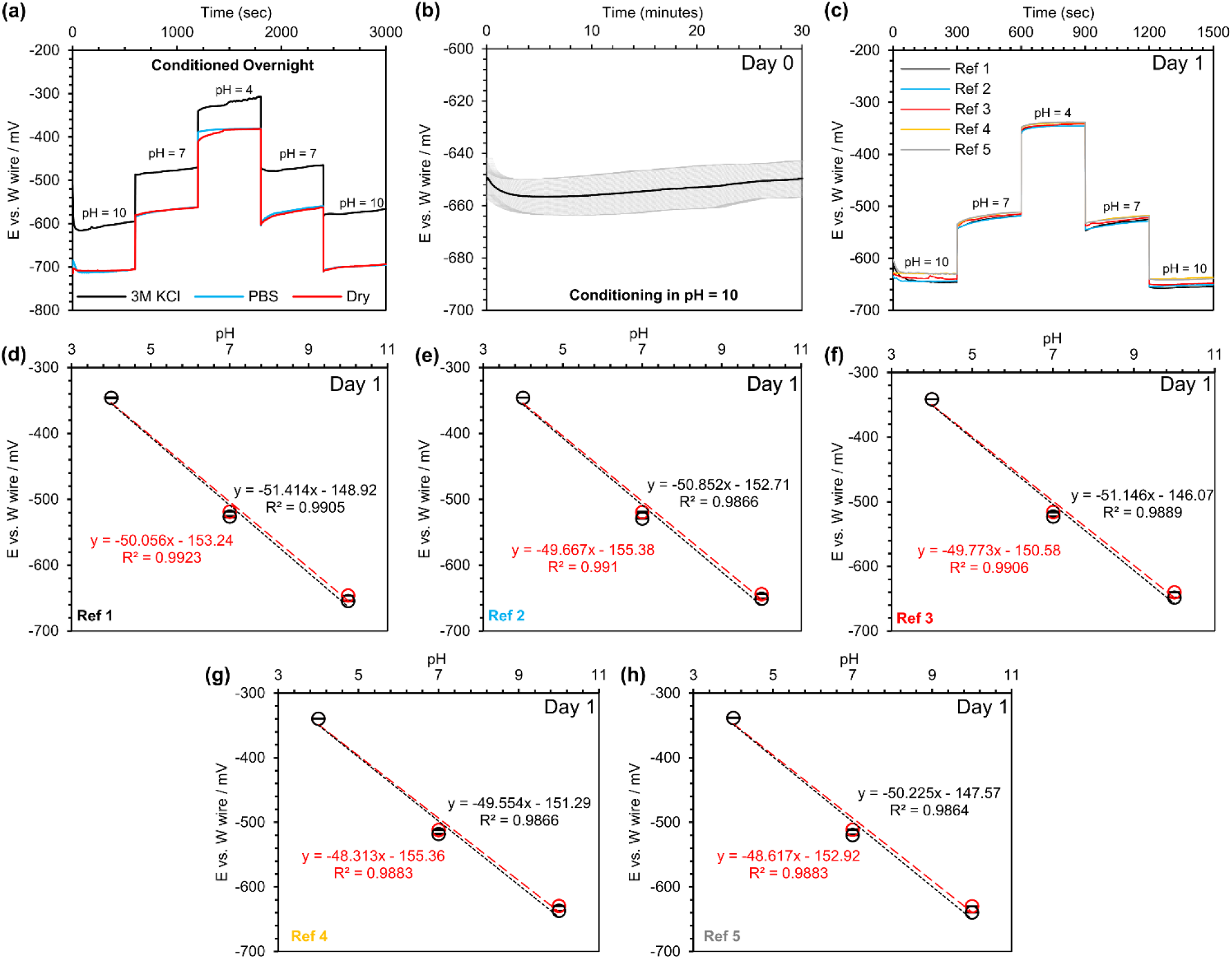
Characterization and testing of tungsten wires with custom-made chlorinated reference electrodes (a) Real-time response of silver wires undergoing 5 minutes of bleached (NaClO) versus a tungsten wire in a series of buffers with the pH of 10, 7, and 4 with a different conditioning treatment, (b) real-time response of 5-minute bleached silver wires versus a tungsten wire in a buffer with the pH of 10 for 30 minutes, (c) real-time response of 5-minute bleached silver wires versus a tungsten wire in standard buffers (pH = 4, 7, and 10) after being conditioned in PBS for one day, (d – h) the calibration curves for each individual sensors. N = 5. The decrease in pH is shown by red colors, and the increase in pH is shown in black color.

The real-time response of custom-made reference electrodes stored in PBS against a tungsten wire was investigated in a standard buffer with a pH of 10 (Figure 3b). The average initial potential was – 650.3 mV, which was consistent with the final average potential (−649.7 mV), showing that a properly-conditioned reference electrode rapidly responded to the new environment. Next, we tested chlorinated reference electrodes versus a tungsten wire. Figure 3c represents the real-time response of these reference electrodes against the tungsten wire after conditioned for one day in PBS, in three different standard buffers (pH = 10, 7, and 4). A smooth transition of the potential electrical signals for all five custom-made reference electrodes was observed when they were transferred from one buffer to another one. However, one of the reference electrodes (Ref 5) had a slightly different potential at a pH of 10 and underwent a larger hysteresis compared to others. Figure 3d to Figure 3h shows the open circuit potential of these reference electrodes at different pH (the calibration curves). The average slope of the calibration curves was – 50 ± 1 mV/pH, which was close to the values reported in other works ^43,44,55^ and was higher compared to commercial reference electrodes reported earlier.

Both custom-made reference electrodes and the tungsten wire were stored in PBS for three more days, and the real-time response of custom-made reference electrodes at day four was evaluated against the tungsten wire under the same conditions (Figure 4a). One of the reference electrodes (Ref 5) failed and did not show significant pH sensitivity. The formed silver/silver chloride layer was fragile and could be damaged due to the movement of the reference electrode. Drift in electrical potentials was observed, most prominent in basic conditions (pH 10), which was 4 – 5 % compared to day 1 (pH = 10). However, the drift in potentials at pH = 4 and pH = 7 were smaller (1 %). The pH response of these reference electrodes against the tungsten wire is shown in Figure 4b to Figure 4e. A decrease in the slope was observed from an average value of ∼ -50.0 mV/pH to -45.7 mV/pH.

**Figure 4:**
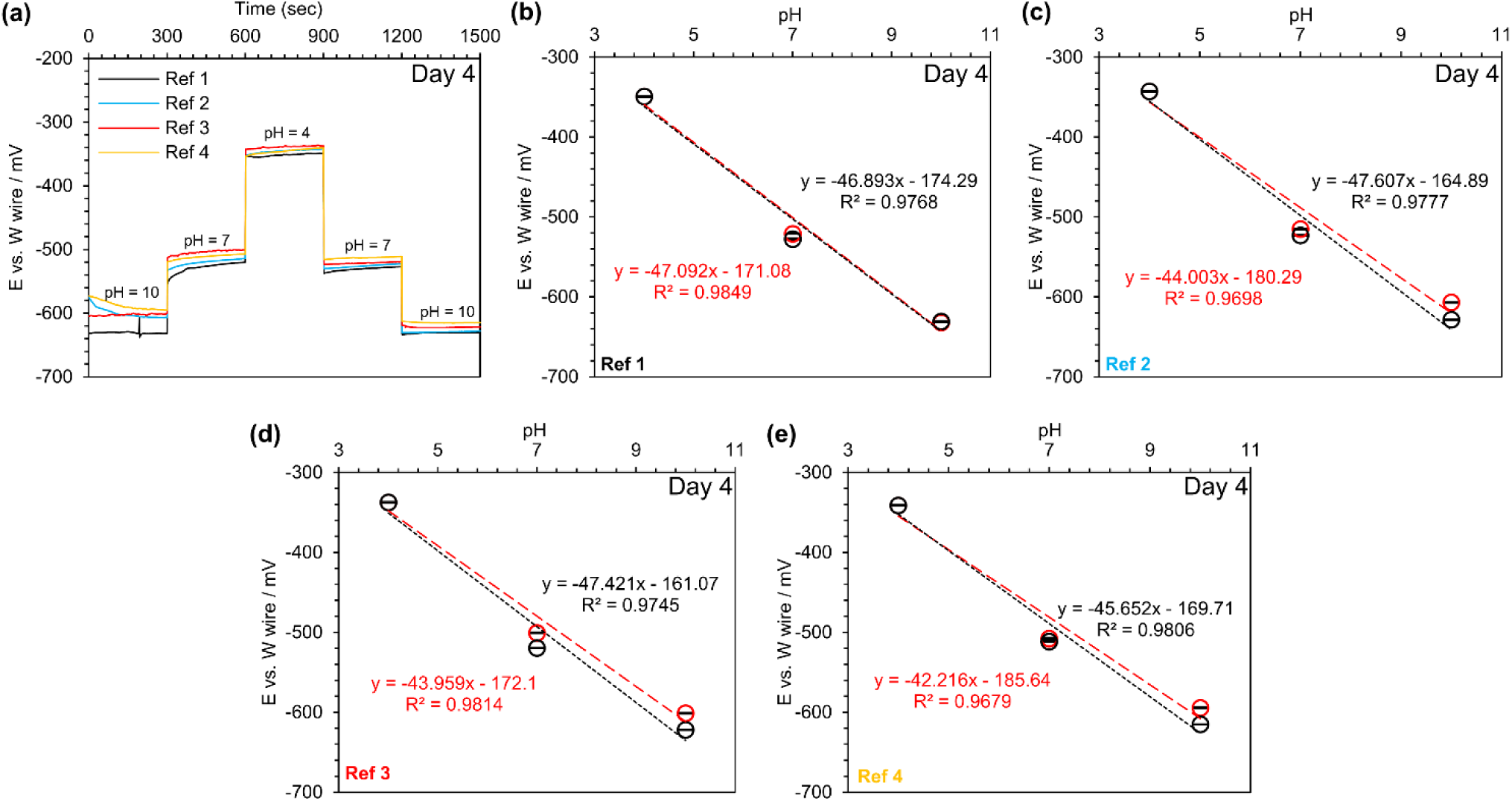
The long-term stability testing of tungsten wires with the custom-made reference electrodes at day 4: (a) real-time response of 5-minute bleached silver wires versus a tungsten wire in standard buffers (pH = 4, 7, and 10) after being stored in PBS for four days, (b – e) the calibration curves for each individual sensors. N = 4. The decrease in pH is shown by red colors, and the increase in pH is shown in black color.

### 4.4 pH sensitivity of tungsten and chlorinated silver wires: Applications in a narrow pH range

To assess the pH sensitivity of tungsten wires with custom-made reference electrodes, different standard buffer solutions with narrower pH values ranging from 7.05 to 9.1 were prepared. In contrast to the previous experiments, where all custom-made reference electrodes (chlorinated silver wires) were tested against a similar tungsten wire to reduce variation in potential measurement, four pairs of pH sensors consisted of tungsten wire and a custom-made reference electrode were fabricated and studied. This would enable us to recognize the difference in pH sensitivity of sensors from one batch to another batch. In addition, the sensors were only conditioned in DI water for an hour to see how fast the sensor could be conditioned. An extra buffer solution with a pH of 7.50 was also added to the experiment when pH was increased from the neutral to the basic conditions. Figure 5a shows the real-time pH response of these sensors. The open circuit potential of these sensors was relatively stable at all pH tested, and a small hysteresis was observed (e.g., E was -650 mV and - 644 mV equal to 1% drift). Figure 5b to Figure 5e represents the open circuit potential of each sensor for the tested pH. While pH was decreased, the average pH sensitivity of the sensors was – 51.1 ± 1.2 mV/pH. However, the average pH sensitivity was decreased to – 47.9 ± 0.3 mV/pH when pH was increased.

**Figure 5:**
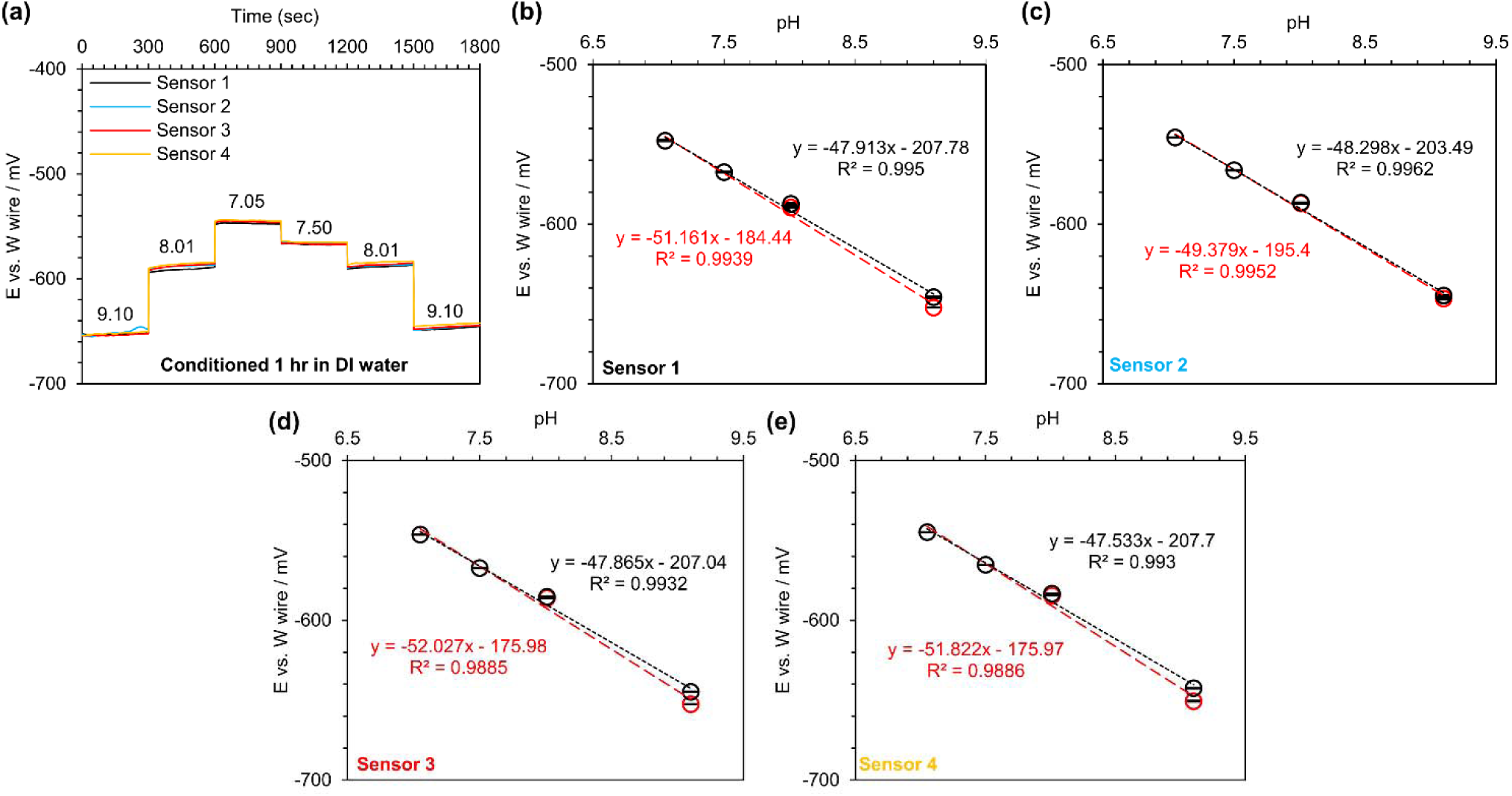
The pH sensitivity of tungsten wires and the custom-made reference electrodes for measuring pH in a narrower range: (a) Real-time response of 5-minute bleached (NaClO) silver wires versus a tungsten wire in a different set of buffers with narrower pH range after being conditioned for 1 hour in DI water, (b – e) the calibration curves for each individual sensors. N = 4. The decrease in pH is shown by red colors, and the increase in pH is shown in black color.

### 4.5 pH performance of sensors in a cell culture medium

PDMS macro-wells with integrated sensors were fabricated and evaluated with cell culture medium. The cell culture medium did not contain bicarbonate as a pH buffer. The macro-well array included three wells, each with a volume of ∼ 800 µL (Figure 6a). In the beginning, wells were rinsed with PBS (pH = 7.4) and the first media (pH = 9.00) for three times. Media was changed with a micropipette, followed by PBS rinse, and rinsing with the next media three times. The real-time pH response of these sensors in media with various pH is plotted in Figure 6b. First, the potential transition between media was smooth and did not show a discernible shift. A slight difference in the open circuit potential between sensor three and two others was seen, which was smaller than 1.5 %. Nonetheless, the pH sensitivity of the sensors was significantly decreased to – 36.4 ± 0.6 mV/pH when they were tested in this cell culture media (Figure 6c to Figure 6e).

**Figure 6:**
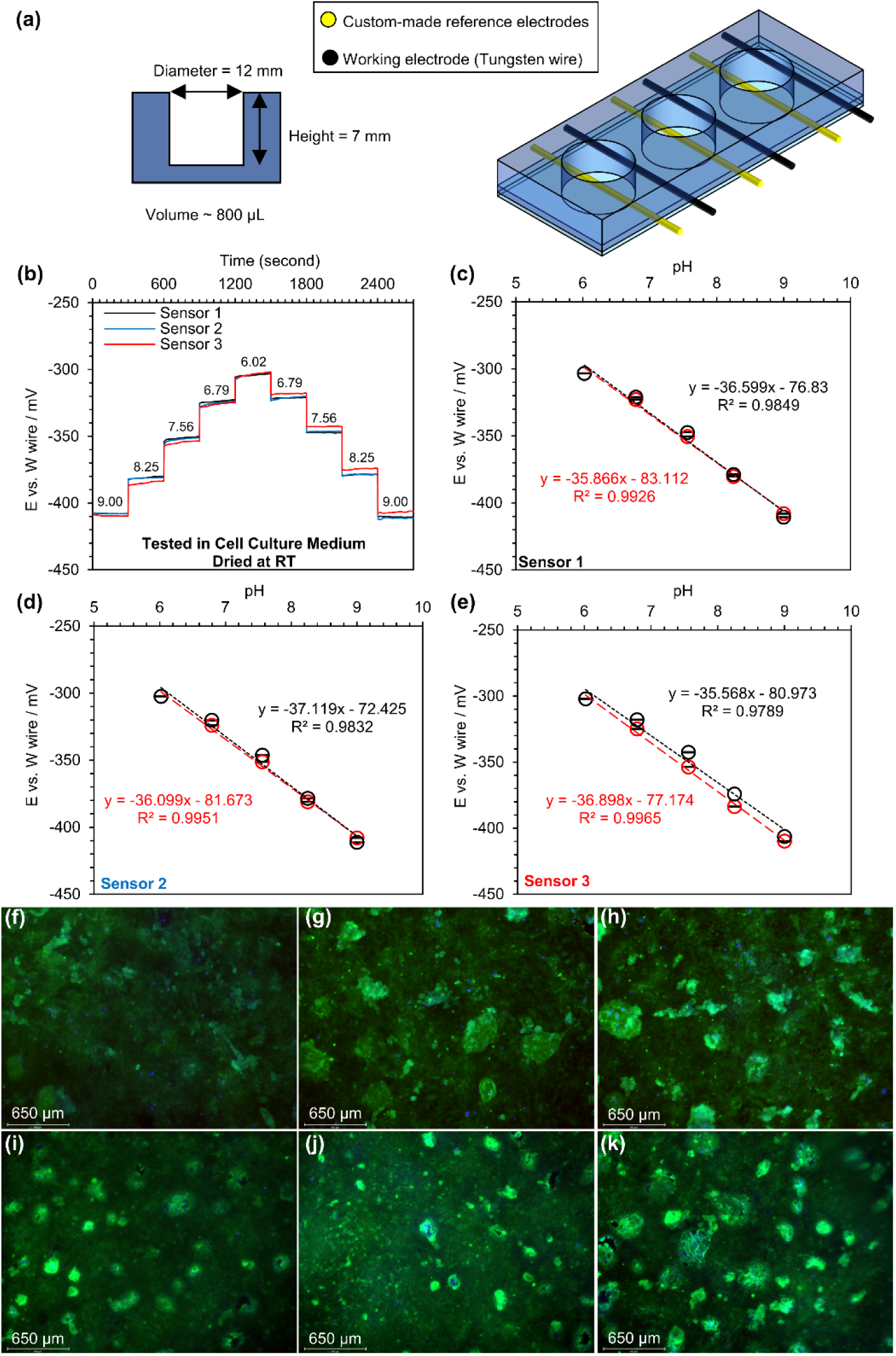
In vitro studies of wire format sensors placed in an array of macro wells: (a) the schematic and dimensions of macro wells, (b) real-time response of 5-minute bleached silver wires versus a tungsten wire in cell culture medium without bicarbonate and adjusted pH after being conditioned in PBS for one day, (c – e) the calibration curves for each individual sensors, (f-h) in vitro viability assay of Calu-3 cells cultured in PDMS macro wells without the present of wire sensors, and (i-k) in vitro viability assay of Calu-3 cells cultured in PDMS macro wells with the present of wire sensors. N = 3. The decrease in pH is shown by red colors, and the increase in pH is shown in black color.

A silver/silver chloride layer without any ion-selective barrier is highly sensitive to chloride ions, resulting in changes to the electrochemical behavior of this type of reference electrode when exposed to a high chloride ion concentration environment ^56–59^. A cell culture media usually contains a significant amount of chloride ions (in order of 100 – 200 mM). Although the custom-made chlorinated reference electrode interfered with chloride ions present in the media, the pH response still behaved linearly and produced a repeatable and predictable response because of having the same chloride concentration in all media.

Calu-3 cells were cultured in three wells with sensor wires and in three wells without wires as a control. In the first 48 hours, the wires were not brought in contact with the media to ensure that all cells uniformly adhered to the collagen-coated PDMS surfaces under identical growth conditions. Then, the media was changed, and its level was increased to cover the wires as well. The cells were grown for 48h following reaching confluency before being stained with Calcein AM and Hoechst dyes for viability assay. Figure 6f to Figure 6k displays the stained cells without observing any negative impacts on cells. In both cases, a uniform distribution of live cells can be seen with some brighter patches of live cells showing that the cells were grown on the top of each other.

Therefore, it can be concluded that the integration of wires (tungsten and a custom-made chlorinated reference electrode) in the wells did not affect the growth and morphology of cells to confluency and even over-confluency.

### 4.6 The effect of pH sensor on cell cytotoxicity or cytokine responses

Lactate dehydrogenase is a cytoplasmic enzyme found in almost all cells in the body, including lung tissue, responsible for converting lactate into pyruvate ^60^. Upon compromise to cell plasma membrane integrity, it is released into the extracellular environment ^60^. As such, it has been used in the past to assess cytotoxicity and cell death ^61^. An LDH Assay was used in our study to quantify the degree of cytotoxicity to Calu-3 cells from the presence of the silver/silver chloride wire and tungsten wire relative to the control condition (no wires). We showed that the presence of the silver/silver chloride + tungsten wires would not induce significant cytotoxicity to the cells. As shown in Figure 7a, LDH concentration as a percent of the maximal LDH release was noticeably lower in the presence and absence of the silver/silver chloride + tungsten wires, respectively. In addition, our data suggested that relative to the maximal LDH release possible, and there was significantly less LDH measured in both the control and silver/silver chloride + tungsten conditions, suggesting that the wires did not induce any toxicity to the cells. Similarly, we showed that the presence of silver/silver chloride +tungsten would not significantly impact the production of IL-6 and IL-8 cytokines, which are associated with inflammation Figure 7b and Figure 7c. Collectively, we demonstrate that cell cytotoxicity and IL-8 and IL-6 responses were not perceivably impacted by our sensor.

**Figure 7:**
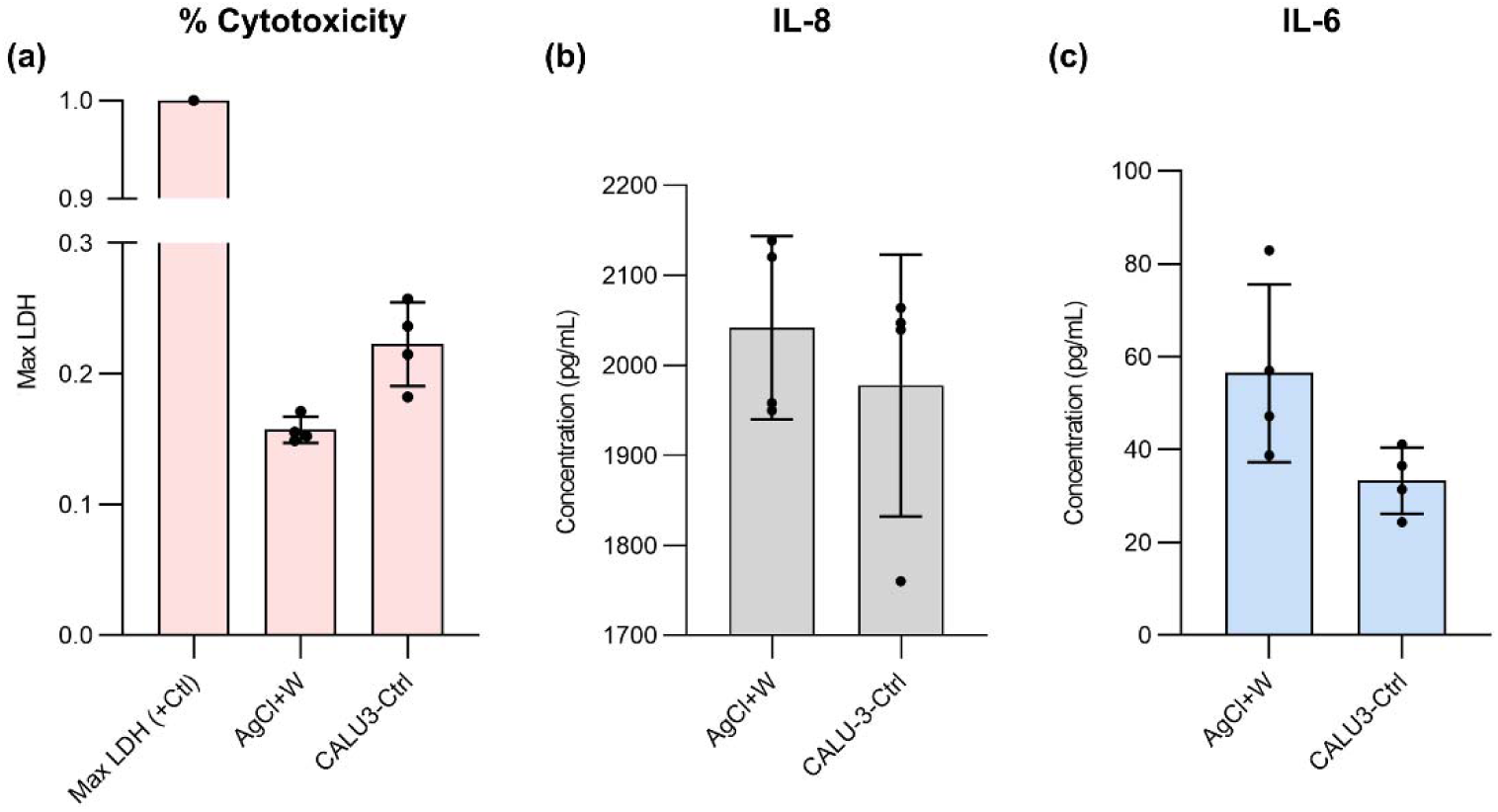
In vitro studies of wire format sensors placed in an array of macro wells: (a) cytotoxicity of wire-format sensors adjacent cells using LDH assay, (b) the production of IL-8, and (c) the production of IL-6. The error bars are standard deviations.

### 4.7 A 3D-printed wire sensors holder for measuring pH inside a Transwell Insert

Commercial pH sensors, even “micro” pH sensors, cannot be introduced into a small reservoir that contains a very small amount of a solution to measure pH in situ. As a result, a 3D-printed holder was designed so that it could introduce both tungsten wire and a custom-made reference electrode onto the cultured cells in a Transwell insert assembled in a new biosensor device - PHAIR, as depicted in Figure 8a. This device could be easily removed form one Transwell insert, rinsed, and placed to another one.

**Figure 8:**
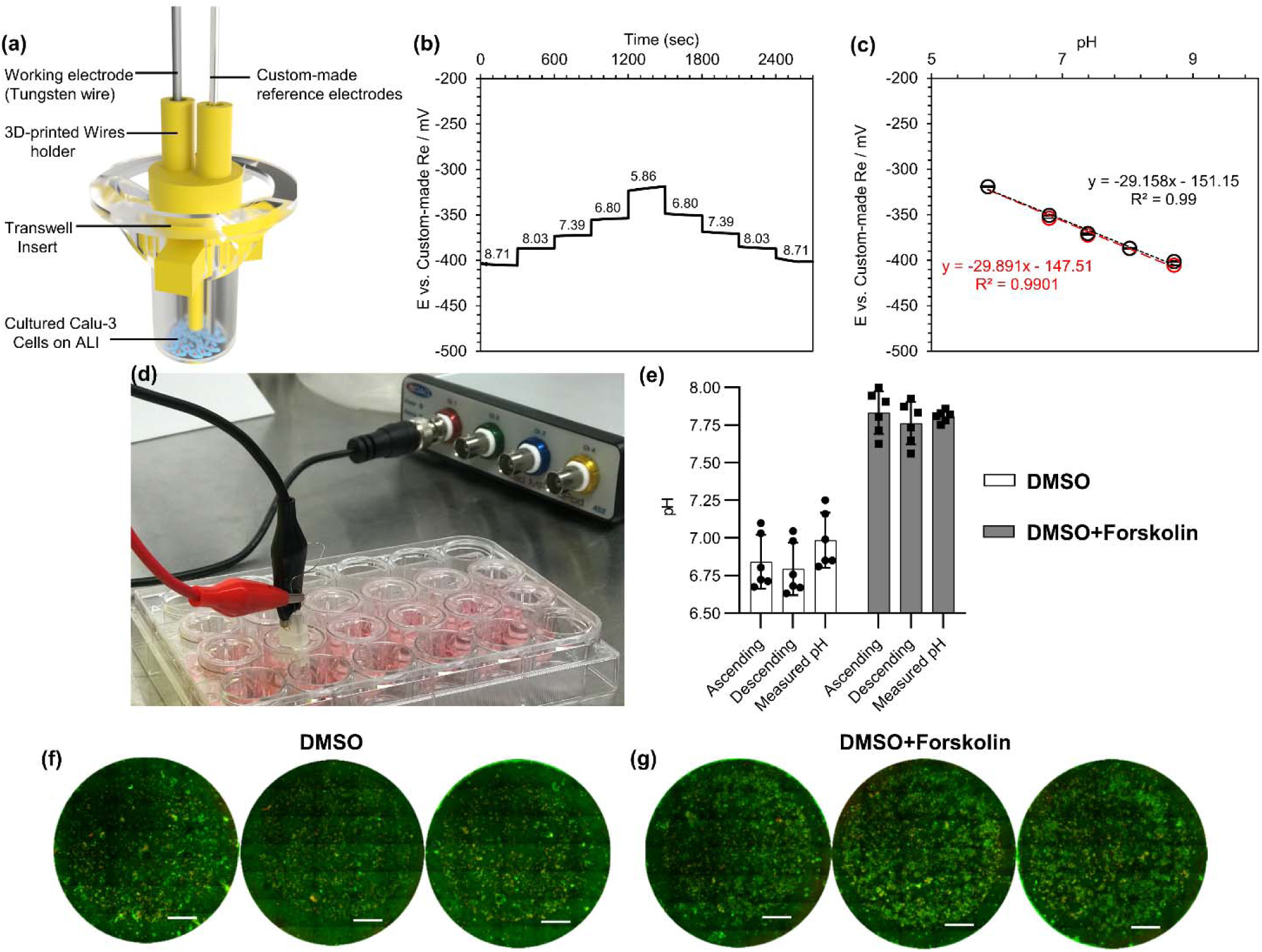
(a) a schematic of the 3D-printed holder for bringing wire sensors in tough with the apical testing solution, (b) real-time response of 5-minute bleached silver wires versus a tungsten wire in home-made Ringer’s solution without any buffer with various pH after being conditioned in PBS for one day, (c) the calibration curve for the sensor (decrease in pH is shown by red colors and increase in pH is shown by black color.), (d) the experimental setup for measuring pH of the apical solution in Transwell insert after being treated by DMSO or DMSO + Forskolin, (e) a comparison between the measured pH by the wire sensor and a commercial pH electrode (apical solutions were collected and pH of them were measured later by the commercial pH meter), and in vitro viability assay of Calu-3 cells treated by (f) DMSO or (g) DMSO + Forskolin. Scale bars are 1 cm.

Calu-3 cells can be grown at an air-liquid interface (ALI) and be used as an *in vitro* model for studying drugs that involve the regulation of cystic fibrosis transmembrane conductance regulator (CFTR) such as cystic fibrosis (CF) ^13,16,62^. Therefore, Calu-3 cells were grown at an ALI and were stimulated by adding forskolin in the basal compartment while a home-made Ringer’s solution was added to the apical side. Since the cells were releasing their secretions to an environment (Ringer’s solution) enriched in chloride ions, the sensor made of a tungsten wire and the custom-made chlorinated reference electrode was tested and calibrated in Ringer’s solutions with various pH prior to the main experiment. The calibrated sensor was thoroughly rinsed with DI water and stored in Ringer’s solution until it was used for measuring the apical pH. The real-time pH response of the sensor in Ringer’s solution with different pH is shown in Figure 8b, suggesting that the open circuit potential was quickly stabilized and experienced a small hysteresis similar to the earlier observations. Nevertheless, a drop in the pH sensitivity of the sensor was observed, which was expected because Ringer’s solution had a high chloride concentration (the pH response was linear and had an R^2^ of ∼ 0.990).

After stimulation of the cells with forskolin, the sensor device was placed in each Transwell, and the pH was reported (Figure 8d). Compared to DMSO vehicle control, 10 µM of forskolin in basal media induced a significant increase in ASL pH according to both methods of pH measurement (Figure 8e). This observation was consistent with current models of ASL pH regulation and results from similar studies, which have reported increases in HCO_3_^-^ secretion, ASL buffer capacity, and pH due to forskolin stimulation ^13,15,16,63,64^. In response to ASL acidification, as simulated in our experiment by the addition of HCO_3_^-^ and K^+^-free saline Ringer’s solution (pH 6.0) to the apical surface, epithelial monolayers are capable of normalizing pH to the alkaline range ^15,17,65,66^. By activating CFTR, a major mechanism of base secretion in Calu-3 cells, through intracellular cAMP elevation, this process can be accelerated. After 3.5 hours without forskolin stimulation, ASL pH, as measured by the commercial microelectrode, was 6.98 ± 0.17 (mean ± SD). With forskolin stimulation, ASL pH was significantly higher at 7.81 ± 0.35, although this effect appears to be greater than what has been reported in the literature, which varies but are typically less than 0.5 ^13,15,17,64^. Such differences may be attributable to heterogeneity in methodology, including cell type, stimulation conditions, and measurement tools. Crucially, the forskolin-induced increase in ASL pH in Figure 8e was verified by two independent methods of measurement; there were no statistical differences between readings from the commercial microelectrode and calculated values from our custom sensors for each experimental condition.

After the *in situ* measurement of pH, the apical solution was collected to check the chloride ion concentration of the solution. The chloride ion concentration in the apical side did not change for both the control samples (DMSO) and the stimulated samples (DMSO + forskolin), confirming that the chloride ion concentration was constant and did not affect the pH evaluation of the samples. After pH measurements, the cells were stained with the live and dead stains to ensure that the cells were alive and not impacted by placing the sensor to the medium (Figure 8f and Figure 8g).

## 5 Conclusion

In this study, we miniaturize pH sensing materials to made micro-wires in the form of main and reference electrodes. Tungsten metallic micro-wire was used as a pH sensing material to measure hydrogen ions. Out of four candidates for costume-made reference electrodes, silver micro-wires with a silver/silver chloride coating showed the most stable response over testing conditions. Both, Tungsten and Silver chloride elected were used to produce a novel device PHAIR to measure pH in simple buffer solutions and in vitro human airway cell culture simultaneously. PHAIR could operate in a smaller volume and cell culture well plates to measure change in pH in close proximity of the cell surface. In our further works, we will explore the usage of PHAIR in more complex cell culture systems and tissues.

## 6 Acknowledgment

This work was supported by a SickKids New Investigator Award. JAH also acknowledges support from the Canada Research Chairs program in Respiratory Mucosal Immunology.

## References

(1) Ghosh, S.; Chang, Y. F.; Yang, D. M.; Chattopadhyay, S. Upconversion Nanoparticle-MOrange Protein FRET Nanoprobes for Self-Ratiometric/Ratiometric Determination of Intracellular PH, and Single Cell PH Imaging. Biosens. Bioelectron. 2020, 155 (December 2019), 112115. https://doi.org/10.1016/j.bios.2020.112115.

(2) Sun, F.; Zhang, P.; Bai, T.; David Galvan, D.; Hung, H. C.; Zhou, N.; Jiang, S.; Yu, Q. Functionalized Plasmonic Nanostructure Arrays for Direct and Accurate Mapping Extracellular PH of Living Cells in Complex Media Using SERS. Biosens. Bioelectron. 2015, 73, 202–207. https://doi.org/10.1016/j.bios.2015.05.060.

(3) Sakata, T.; Sugimoto, H.; Saito, A. Live Monitoring of Microenvironmental PH Based on Extracellular Acidosis around Cancer Cells with Cell-Coupled Gate Ion-Sensitive Field-Effect Transistor. Anal. Chem. 2018, 90 (21), 12731–12736. https://doi.org/10.1021/acs.analchem.8b03070.

(4) McBeth, C.; Dughaishi, R. Al; Paterson, A.; Sharp, D. Ubiquinone Modified Printed Carbon Electrodes for Cell Culture PH Monitoring. Biosens. Bioelectron. 2018, 113 (February), 46–51. https://doi.org/10.1016/j.bios.2018.04.052.

(5) Ges, I. A.; Ivanov, B. L.; Schaffer, D. K.; Lima, E. A.; Werdich, A. A.; Baudenbacher, F. J. Thin-Film IrOxpH Microelectrode for Microfluidic-Based Microsystems. Biosens. Bioelectron. 2005, 21 (2), 248–256. https://doi.org/10.1016/j.bios.2004.09.021.

(6) Weltin, A.; Slotwinski, K.; Kieninger, J.; Moser, I.; Jobst, G.; Wego, M.; Ehret, R.; Urban, G. A. Cell Culture Monitoring for Drug Screening and Cancer Research: A Transparent, Microfluidic, Multi-Sensor Microsystem. Lab Chip 2014, 14 (1), 138–146. https://doi.org/10.1039/c3lc50759a.

(7) Hirota, J. A.; Knight, D. A. Human Airway Epithelial Cell Innate Immunity: Relevance to Asthma. Curr. Opin. Immunol. 2012, 24 (6), 740–746. https://doi.org/10.1016/j.coi.2012.08.012.

(8) Haq, I. J.; Gray, M. A.; Garnett, J. P.; Ward, C.; Brodlie, M. Airway Surface Liquid Homeostasis in Cystic Fibrosis: Pathophysiology and Therapeutic Targets. Thorax 2016, 71 (3), 284–287. https://doi.org/10.1136/thoraxjnl-2015-207588.

(9) Fischer, H.; Widdicombe, J. H. Mechanisms of Acid and Base Secretion by the Airway Epithelium. J. Membr. Biol. 2006, 211 (3), 139–150. https://doi.org/10.1007/s00232-006-0861-0.

(10) Ricciardolo, F. L. M.; Gaston, B.; Hunt, J. Acid Stress in the Pathology of Asthma. J. Allergy Clin. Immunol. 2004, 113 (4), 610–619. https://doi.org/10.1016/j.jaci.2003.12.034.

(11) Ostedgaard, L. S.; Baldursson, O.; Welsh, M. J. Regulation of the Cystic Fibrosis Transmembrane Conductance Regulator Cl- Channel by Its R Domain. J. Biol. Chem. 2001, 276 (11), 7689–7692. https://doi.org/10.1074/jbc.R100001200.

(12) Tang, L.; Fatehi, M.; Linsdell, P. Mechanism of Direct Bicarbonate Transport by the CFTR Anion Channel. J. Cyst. Fibros. 2009, 8 (2), 115–121. https://doi.org/10.1016/j.jcf.2008.10.004.

(13) Huang, J.; Kim, D.; Shan, J.; Abu-Arish, A.; Luo, Y.; Hanrahan, J. W. Most Bicarbonate Secretion by Calu-3 Cells Is Mediated by CFTR and Independent of Pendrin. Physiol. Rep. 2018, 6 (5), 1–17. https://doi.org/10.14814/phy2.13641.

(14) Simonin, J.; Bille, E.; Crambert, G.; Noel, S.; Dreano, E.; Edwards, A.; Hatton, A.; Pranke, I.; Villeret, B.; Cottart, C. H.; Vrel, J. P.; Urbach, V.; Baatallah, N.; Hinzpeter, A.; Golec, A.; Touqui, L.; Nassif, X.; Galietta, L. J. V.; Planelles, G.; Sallenave, J. M.; Edelman, A.; Sermet-Gaudelus, I. Airway Surface Liquid Acidification Initiates Host Defense Abnormalities in Cystic Fibrosis. Sci. Rep. 2019, 9 (1), 1–11. https://doi.org/10.1038/s41598-019-42751-4.

(15) Coakley, R. D.; Grubb, B. R.; Paradiso, A. M.; Gatzy, J. T.; Johnson, L. G.; Kreda, S. M.; O’Neal, W. K.; Boucher, R. C. Abnormal Surface Liquid PH Regulation by Cultured Cystic Fibrosis Bronchial Epithelium. Proc. Natl. Acad. Sci. U. S. A. 2003, 100 (26), 16083–16088. https://doi.org/10.1073/pnas.2634339100.

(16) Kim, D.; Liao, J.; Hanrahan, J. W. The Buffer Capacity of Airway Epithelial Secretions. Front. Physiol. 2014, 5 JUN (June), 1–11. https://doi.org/10.3389/fphys.2014.00188.

(17) Garnett, J. P.; Kalsi, K. K.; Sobotta, M.; Bearham, J.; Carr, G.; Powell, J.; Brodlie, M.; Ward, C.; Tarran, R.; Baines, D. L. Hyperglycaemia and Pseudomonas Aeruginosa Acidify Cystic Fibrosis Airway Surface Liquid by Elevating Epithelial Monocarboxylate Transporter 2 Dependent Lactate-H + Secretion. Sci. Rep. 2016, 6 (November), 1–13. https://doi.org/10.1038/srep37955.

(18) Berkebile, A. R.; Bartlett, J. A.; Alaiwa, M. A.; Varga, S. M.; Power, U. F.; McCray, P. B. Airway Surface Liquid Has Innate Antiviral Activity That Is Reduced in Cystic Fibrosis. Am. J. Respir. Cell Mol. Biol. 2020, 62 (1), 104–111. https://doi.org/10.1165/rcmb.2018-0304OC.

(19) Pezzulo, A. A.; Tang, X. X.; Hoegger, M. J.; Abou Alaiwa, M. H., Ramachandran, S.; Moninger, T. O.; Karp, P. H.; Wohlford-Lenane, C. L.; Haagsman, H. P.; Eijk, M. Van; Bánfi, B.; Horswill, A. R.; Stoltz, D. A.; Mc Cray, P. B., Welsh, M. J.; Zabner, J. Reduced Airway Surface PH Impairs Bacterial Killing in the Porcine Cystic Fibrosis Lung. Nature 2012, 487 (7405), 109–113. https://doi.org/10.1038/nature11130.

(20) Gianotti, A.; Capurro, V.; Delpiano, L.; Mielczarek, M.; García Lvalverde, M., CarreiraLbarral, I.; Ludovico, A.; Fiore, M.; Baroni, D.; Moran, O.; Quesada, R.; Caci, E. Small Molecule Anion Carriers Correct Abnormal Airway Surface Liquid Properties in Cystic Fibrosis Airway Epithelia. Int. J. Mol. Sci. 2020, 21 (4), 1–14. https://doi.org/10.3390/ijms21041488.

(21) Delpiano, L.; Thomas, J. J.; Yates, A. R.; Rice, S. J.; Gray, M. A.; Saint-Criq, V. Esomeprazole Increases Airway Surface Liquid Ph in Primary Cystic Fibrosis Epithelial Cells. Front. Pharmacol. 2018, 9 (December), 1–15. https://doi.org/10.3389/fphar.2018.01462.

(22) Boj, S. F.; Vonk, A. M.; Statia, M.; Su, J.; Vries, R. R. G.; Beekman, J. M.; Clevers, H. Forskolin-Induced Swelling in Intestinal Organoids: An in Vitro Assay for Assessing Drug Response in Cystic Fibrosis Patients. J. Vis. Exp. 2017, e55159 (120), 1–12. https://doi.org/10.3791/55159.

(23) Schnúr, A.; Premchandar, A.; Bagdany, M.; Lukacs, G. L. Phosphorylation-Dependent Modulation of CFTR Macromolecular Signalling Complex Activity by Cigarette Smoke Condensate in Airway Epithelia. Sci. Rep. 2019, 9 (1), 1–19. https://doi.org/10.1038/s41598-019-48971-y.

(24) Li, C.; Dandridge, K. S.; Di, A.; Marrs, K. L.; Harris, E. L.; Roy, K.; Jackson, J. S.; Makarova, N. V.; Fujiwara, Y.; Farrar, P. L.; Nelson, D. J.; Tigyi, G. J.; Naren, A. P. Lysophosphatidic Acid Inhibits Cholera Toxin-Induced Secretory Diarrhea through CFTR-Dependent Protein Interactions. J. Exp. Med. 2005, 202 (7), 975–986. https://doi.org/10.1084/jem.20050421.

(25) Gorziglia, M.; Hoshino, Y.; Buckler-White, A.; Blumentals, I.; Glass, R.; Flores, J.; Kapikian, A. Z.; Chanock, R. M. Conservation of Amino Acid Sequence of VP8 and Cleavage Region of 84-KDa Outer Capsid Protein among Rotaviruses Recovered from Asymptomatic Neonatal Infection. Proc. Natl. Acad. Sci. U. S. A. 1986, 83 (18), 7039–7043. https://doi.org/10.1073/pnas.83.18.7039.

(26) Schultz, A.; Puvvadi, R.; Borisov, S. M.; Shaw, N. C.; Klimant, I.; Berry, L. J.; Montgomery, S. T.; Nguyen, T.; Kreda, S. M.; Kicic, A.; Noble, P. B.; Button, B.; Stick, S. M. Airway Surface Liquid PH Is Not Acidic in Children with Cystic Fibrosis. Nat. Commun. 2017, 8 (1), 1–8. https://doi.org/10.1038/s41467-017-00532-5.

(27) Dang, W.; Manjakkal, L.; Navaraj, W. T.; Lorenzelli, L.; Vinciguerra, V.; Dahiya, R. Stretchable Wireless System for Sweat PH Monitoring. Biosens. Bioelectron. 2018, 107 (December 2017), 192–202. https://doi.org/10.1016/j.bios.2018.02.025.

(28) Kajisa, T.; Yanagimoto, Y.; Saito, A.; Sakata, T. Biocompatible Poly(Catecholamine)-Film Electrode for Potentiometric Cell Sensing. ACS Sensors 2018, 3 (2), 476–483. https://doi.org/10.1021/acssensors.7b00897.

(29) Yoon, J. H.; Kim, S. M.; Park, H. J.; Kim, Y. K.; Oh, D. X.; Cho, H. W.; Lee, K. G.; Hwang, S. Y.; Park, J.; Choi, B. G. Highly Self-Healable and Flexible Cable-Type PH Sensors for Real-Time Monitoring of Human Fluids. Biosens. Bioelectron. 2020, 150 (November 2019), 111946. https://doi.org/10.1016/j.bios.2019.111946.

(30) Liu, M. M.; Guo, Z. Z.; Liu, H.; Li, S. H.; Chen, Y.; Zhong, Y.; Lei, Y.; Lin, X. H.; Liu, A. L. Paper-Based 3D Culture Device Integrated with Electrochemical Sensor for the on-Line Cell Viability Evaluation of Amyloid-Beta Peptide Induced Damage in PC12 cells. Biosens. Bioelectron. 2019, 144 (August), 111686. https://doi.org/10.1016/j.bios.2019.111686.

(31) Bao, B.; Yang, Z.; Liu, Y.; Xu, Y.; Gu, B.; Chen, J.; Su, P.; Tong, L.; Wang, L. Two-Photon Semiconducting Polymer Nanoparticles as a New Platform for Imaging of Intracellular PH Variation. Biosens. Bioelectron. 2019, 126 (October 2018), 129–135. https://doi.org/10.1016/j.bios.2018.10.027.

(32) Ges, I. A.; Ivanov, B. L.; Werdich, A. A.; Baudenbacher, F. J. Differential PH Measurements of Metabolic Cellular Activity in Nl Culture Volumes Using Microfabricated Iridium Oxide Electrodes. Biosens. Bioelectron. 2007, 22 (7), 1303–1310. https://doi.org/10.1016/j.bios.2006.05.033.

(33) Muñoz-Berbel, X.; Rodríguez-Rodríguez, R., Vigués, N.; Demming, S.; Mas, J.; Büttgenbach, S.; Verpoorte, E.; Ortiz, P.; Llobera, A. Monolithically Integrated Biophotonic Lab-on-a-Chip for Cell Culture and Simultaneous PH Monitoring. Lab Chip 2013, 13 (21), 4239–4247. https://doi.org/10.1039/c3lc50746g.

(34) Magnusson, E. B.; Halldorsson, S.; Fleming, R. M. T.; Leosson, K. Real-Time Optical PH Measurement in a Standard Microfluidic Cell Culture System. Biomed. Opt. Express 2013, 4 (9), 1749. https://doi.org/10.1364/boe.4.001749.

(35) Shaegh, S. A. M.; De Ferrari, F.; Zhang, Y. S.; Nabavinia, M.; Mohammad, N. B.; Ryan, J.; Pourmand, A.; Laukaitis, E.; Sadeghian, R. B.; Nadhman, A.; Shin, S. R.; Nezhad, A. S.; Khademhosseini, A.; Dokmeci, M. R. A Microfluidic Optical Platform for Real-Time Monitoring of PH and Oxygen in Microfluidic Bioreactors and Organ-on-Chip Devices. Biomicrofluidics 2016, 10 (4), 1–14. https://doi.org/10.1063/1.4955155.

(36) Ghoneim, M. T.; Nguyen, A.; Dereje, N.; Huang, J.; Moore, G. C.; Murzynowski, P. J.; Dagdeviren, C. Recent Progress in Electrochemical PH-Sensing Materials and Configurations for Biomedical Applications. Chem. Rev. 2019, 119 (8), 5248–5297. https://doi.org/10.1021/acs.chemrev.8b00655.

(37) Yao, S.; Wang, M.; Madou, M. A PH Electrode Based on Melt-Oxidized Iridium Oxide. J. Electrochem. Soc. 2001, 148 (4), H29. https://doi.org/10.1149/1.1353582.

(38) Huang, X. R.; Ren, Q. Q.; Yuan, X. J.; Wen, W.; Chen, W.; Zhan, D. P. Iridium Oxide Based Coaxial PH Ultramicroelectrode. Electrochem. commun. 2014, 40, 35–37. https://doi.org/10.1016/j.elecom.2013.12.012.

(39) Mingels, R. H. G.; Kalsi, S.; Cheong, Y.; Morgan, H. Iridium and Ruthenium Oxide Miniature PH Sensors: Long-Term Performance. Sensors Actuators, B Chem. 2019, 297 (June), 126779. https://doi.org/10.1016/j.snb.2019.126779.

(40) Kim, T. Y.; Yang, S. Fabrication Method and Characterization of Electrodeposited and Heat-Treated Iridium Oxide Films for PH Sensing. Sensors Actuators, B Chem. 2014, 196, 31–38. https://doi.org/10.1016/j.snb.2014.02.004.

(41) Warren, A.; Nylund, A.; Olefjord, I. Oxidation of Tungsten and Tungsten Carbide in Dry and Humid Atmospheres. Int. J. Refract. Met. Hard Mater. 1996, 14 (5–6), 345–353. https://doi.org/10.1016/s0263-4368(96)00027-3.

(42) Anik, M.; Osseo-Asare, K. Effect of PH on the Anodic Behavior of Tungsten. J. Electrochem. Soc. 2002, 149 (6), B224. https://doi.org/10.1149/1.1471544.

(43) Guo, Q.; Wu, X.; Han, E. H.; Ke, W. PH Response Behaviors and Mechanisms of Different Tungsten/Tungsten Oxide Electrodes for Long-Term Monitoring. J. Electroanal. Chem. 2016, 782, 91–97. https://doi.org/10.1016/j.jelechem.2016.10.026.

(44) Wen, Y.; Wang, X. Characterization and Application of a Metallic Tungsten Electrode for Potentiometric PH Measurements. J. Electroanal. Chem. 2014, 14–715, 45–50. https://doi.org/10.1016/j.jelechem.2013.12.031.

(45) Hussain, M.; Ibupoto, Z. H.; Abbasi, M. A.; Nur, O.; Willander, M. Effect of Anions on the Morphology of Co3O4 Nanostructures Grown by Hydrothermal Method and Their PH Sensing Application. J. Electroanal. Chem. 2014, 717–718, 78–82. https://doi.org/10.1016/j.jelechem.2014.01.011.

(46) Scandurra, A.; Bruno, E.; Condorelli, G. G.; Grimaldi, M. G.; Mirabella, S. Microscopic Model for PH Sensing Mechanism in Zinc-Based Nanowalls. Sensors Actuators, B Chem. 2019, 296 (January), 126614. https://doi.org/10.1016/j.snb.2019.05.091.

(47) Lue, C. E.; Wang, I. S.; Huang, C. H.; Shiao, Y. T.; Wang, H. C.; Yang, C. M.; Hsu, S. H.; Chang, C. Y.; Wang, W.; Lai, C. S. PH Sensing Reliability of Flexible ITO/PET Electrodes on EGFETs Prepared by a Roll-to-Roll Process. Microelectron. Reliab. 2012, 52 (8), 1651–1654. https://doi.org/10.1016/j.microrel.2011.10.026.

(48) Tsai, C. N.; Chou, J. C.; Sun, T. P.; Hsiung, S. K. Study on the Sensing Characteristics and Hysteresis Effect of the Tin Oxide PH Electrode. Sensors Actuators, B Chem. 2005, 108 (1-2 SPEC. ISS.), 877–882. https://doi.org/10.1016/j.snb.2004.11.050.

(49) Yin, L. Te; Chou; J. C., Chung, W. Y.; Sun, T. P.; Hsiung, S. K. Study of Indium Tin Oxide Thin Film for Separative Extended Gate ISFET. Mater. Chem. Phys. 2001, 70 (1), 12–16. https://doi.org/10.1016/S0254-0584(00)00373-4.

(50) Qin, Y.; Alam, A. U.; Pan, S.; Howlader, M. M. R.; Ghosh, R.; Selvaganapathy, P. R.; Wu, Y.; Deen, M. J. Low-Temperature Solution Processing of Palladium/Palladium Oxide Films and Their PH Sensing Performance. Talanta 2016, 146, 517–524. https://doi.org/10.1016/j.talanta.2015.08.062.

(51) Qin, Y.; Alam, A. U.; Howlader, M. M. R.; Hu, N. X.; Deen, M. J. Inkjet Printing of a Highly Loaded Palladium Ink for Integrated, Low-Cost PH Sensors. Adv. Funct. Mater. 2016, 26 (27), 4923–4933. https://doi.org/10.1002/adfm.201600657.

(52) Das, A.; Ko, D. H.; Chen, C. H.; Chang, L. B.; Lai, C. S.; Chu, F. C.; Chow, L.; Lin, R. M. Highly Sensitive Palladium Oxide Thin Film Extended Gate FETs as PH Sensor. Sensors Actuators, B Chem. 2014, 205, 199–205. https://doi.org/10.1016/j.snb.2014.08.057.

(53) Grdeń, M.; Łukaszewski, M.; Jerkiewicz, G.; Czerwiński, A. Electrochemical Behaviour of Palladium Electrode: Oxidation, Electrodissolution and Ionic Adsorption. Electrochimica Acta. 2008, pp 7583–7598. https://doi.org/10.1016/j.electacta.2008.05.046.

(54) Barlag, R.; Nyasulu, F.; Starr, R.; Silverman, J.; Arthasery, P.; McMills, L. A Student-Made Silver-Silver Chloride Reference Electrode for the General Chemistry Laboratory: ∼10 Min Preparation. J. Chem. Educ. 2014, 91 (5), 766–768. https://doi.org/10.1021/ed400722e.

(55) Pašti, I. A.; Lazarević-Pašti, T.; Mentus, S. V. Switching between Voltammetry and Potentiometry in Order to Determine H+ or OH- Ion Concentration over the Entire PH Scale by Means of Tungsten Disk Electrode. J. Electroanal. Chem. 2012, 665, 83–89. https://doi.org/10.1016/j.jelechem.2011.11.019.

(56) Ke, X. Micro-Fabricated Electrochemical Chloride Ion Sensors: From the Present to the Future. Talanta 2020, 211 (October 2019), 120734. https://doi.org/10.1016/j.talanta.2020.120734.

(57) Choi, D. H.; Thaxton, A.; Jeong, I. cheol; Kim, K.; Sosnay, P. R.; Cutting, G. R.; Searson, P. C. Sweat Test for Cystic Fibrosis: Wearable Sweat Sensor vs. Standard Laboratory Test. J. Cyst. Fibros. 2018, 17 (4), e35.–e38. https://doi.org/10.1016/j.jcf.2018.03.005.

(58) Choi, D. H.; Li, Y.; Cutting, G. R.; Searson, P. C. A Wearable Potentiometric Sensor with Integrated Salt Bridge for Sweat Chloride Measurement. Sensors Actuators, B Chem. 2017, 250, 673–678. https://doi.org/10.1016/j.snb.2017.04.129.

(59) Mousavi, M. P. S.; Ainla, A.; Tan, E. K. W.; Abd El-Rahman, M., Yoshida, Y.; Yuan, L.; Sigurslid, H. H.; Arkan, N.; Yip, M. C.; Abrahamsson, C. K.; Homer-Vanniasinkam, S.; Whitesides, G. M. Ion Sensing with Thread-Based Potentiometric Electrodes. Lab Chip 2018, 18 (15), 2279–2290. https://doi.org/10.1039/c8lc00352a.

(60) Kumar, P.; Nagarajan, A.; Uchil, P. D. Analysis of Cell Viability by the Lactate Dehydrogenase Assay. Cold Spring Harb. Protoc. 2018, 2018 (6), 465–468. https://doi.org/10.1101/pdb.prot095497.

(61) Smith, S. M.; Wunder, M. B.; Norris, D. A.; Shellman, Y. G. A Simple Protocol for Using a LDH-Based Cytotoxicity Assay to Assess the Effects of Death and Growth Inhibition at the Same Time. PLoS One 2011, 6 (11). https://doi.org/10.1371/journal.pone.0026908.

(62) Zhu, Y.; Chidekel, A.; Shaffer, T. H. Cultured Human Airway Epithelial Cells (Calu-3): A Model of Human Respiratory Function, Structure, and Inflammatory Responses. Crit. Care Res. Pract. 2010, 2010, 1–8. https://doi.org/10.1155/2010/394578.

(63) Shamsuddin, A. K. M.; Quinton, P. M. Native Small Airways Secrete Bicarbonate. Am. J. Respir. Cell Mol. Biol. 2014, 50 (4), 796–804. https://doi.org/10.1165/rcmb.2013-0418OC.

(64) Tang, X. X.; Stoltz, D. A.; Welsh, M. J.; Tang, X. X.; Ostedgaard, L. S.; Hoegger, M. J.; Moninger, T. O.; Karp, P. H.; Mcmenimen, J. D.; Choudhury, B.; Varki, A.; Stoltz, D. A.; Welsh, M. J. Acidic PH Increases Airway Surface Liquid Viscosity in Cystic Fibrosis. J. Clin. Invest. 2016, 126 (3), 879–891. https://doi.org/10.1172/JCI83922.Mucociliary.

(65) Jayaraman, S.; Song, Y.; Verkman, A. S. Airway Surface Liquid PH in Well-Differentiated Airway Epithelial Cell Cultures and Mouse Trachea. Am. J. Physiol. - Cell Physiol. 2001, 281 (5 50-5), 1504–1511. https://doi.org/10.1152/ajpcell.2001.281.5.c1504.

(66) Krouse, M. E.; Talbott, J. F.; Lee, M. M.; Nam, S. J.; Wine, J. J. Acid and Base Secretion in the Calu-3 Model of Human Serous Cells. Am. J. Physiol. - Lung Cell. Mol. Physiol. 2004, 287 (6 31-6), 1274–1283. https://doi.org/10.1152/ajplung.00036.2004.

